# Epithelial invagination by vertical telescoping

**DOI:** 10.1101/515981

**Authors:** Jingjing Li, Andrew D. Economou, Jeremy B.A. Green

## Abstract

Epithelial bending is a fundamental process that shapes organs during development. All currently known mechanisms involve cells locally changing shape from columnar to wedge-shaped. Often this shape change occurs by cytoskeletal contraction at cell apices (“apical constriction”) but mechanisms such as basal nuclear positioning (“basal wedging”) or extrinsic compression are also known. Here we demonstrate a completely different mechanism which occurs without cell wedging. In mammalian salivary glands and teeth, we show that initial invagination occurs through coordinated vertical cell movement. Specifically, we show that cells towards the periphery of the placode move vertically upwards while their more central neighbours move downwards to create the invagination. We further show that this occurs by active cell-on-cell migration: outer cells migrate with an apical leading edge protrusion, depressing the central cells to “telescope” the epithelium downwards into the underlaying mesenchyme. Cells remain basally attached to the underlying lamina while their apical protrusions are dynamic and planar polarised centripetally. These protrusions depend on the actin cytoskeleton, and inhibition of the branching molecule Arp2/3 inhibits them and the invagination. FGF and Hedgehog morphogen signals are also required, with FGF providing a directional cue. These findings show that epithelial bending can be achieved by novel morphogenetic mechanism of coordinated cell rearrangement quite distinct from previously recognised invagination processes.

## Main Text

Ectodermal organs, including hair follicles, teeth, mammary glands, and smaller glands such as mucous, salivary, sweat, lacrimal and sebaceous, are all regulated by a common set of genes, all form from a placode (epithelial thickening) and are all affected simultaneously by congenital ectodermal dysplasias ^1, 2^. We previously showed that in mammalian tooth germs, mammary glands and hair follicles, the epithelium invaginates when it stratifies and cells in the suprabasal layers (suprabasal cells) migrate centripetally and pull radially inwards on a ring of underlying basal layer cells, both bending the epithelium and narrowing the “neck” of the bud (Fig. 1a-c). In this mechanism, canopy contraction, basal layer cells become wedge-shaped due to the extrinsic contraction of the overlying canopy ^3^. Canopy contraction, however, only explains invagination in a solid organ primordium as is seen in so-called “bud” or “peg” stages of teeth, hair follicles and mammary glands. By contrast, salivary glands (SGs), as well as the other glandular ectodermal organs, initially invaginate with a hollow space and no canopy above the invagination (Fig. 1d-f). Close inspection of early tooth invagination shows that before contractile canopy formation, it too initially invaginates as a hollow shape (Fig. 1a). The absence of a canopy rules out canopy contraction, but raises the question: how does this hollow invagination occur? We tested for apical constriction in SG placodes by staining for its hallmarks, namely apical enrichment of F-actin and diphosphomyosin light chain-2 (ref. ^4^). We found that neither F-actin nor diphosphomyosin was detectably enriched in invaginating salivary germs compared to nearby flat oral epithelium (Fig. 1g-j), nor did we see circumferential actin cables which are seen in some other invaginating systems ^5, 6^ (data not shown). Upon cell shape analysis, using mosaic labelling of individual cells in the epithelium, we unexpectedly found no significantly apically narrowed, wedge-shaped cells in any region of the invaginating SG placode, with no significant deviation from columnar shape (Fig.1k-n). This immediately ruled out not only apical constriction but also some other alternative epithelial bending mechanisms, including basal relaxation and basal wedging ^7–13^. Consistent with this, nuclear height was uniformly in the middle of the apicobasal axis throughout the invaginating placode (Fig.1o).

**Fig. 1.**
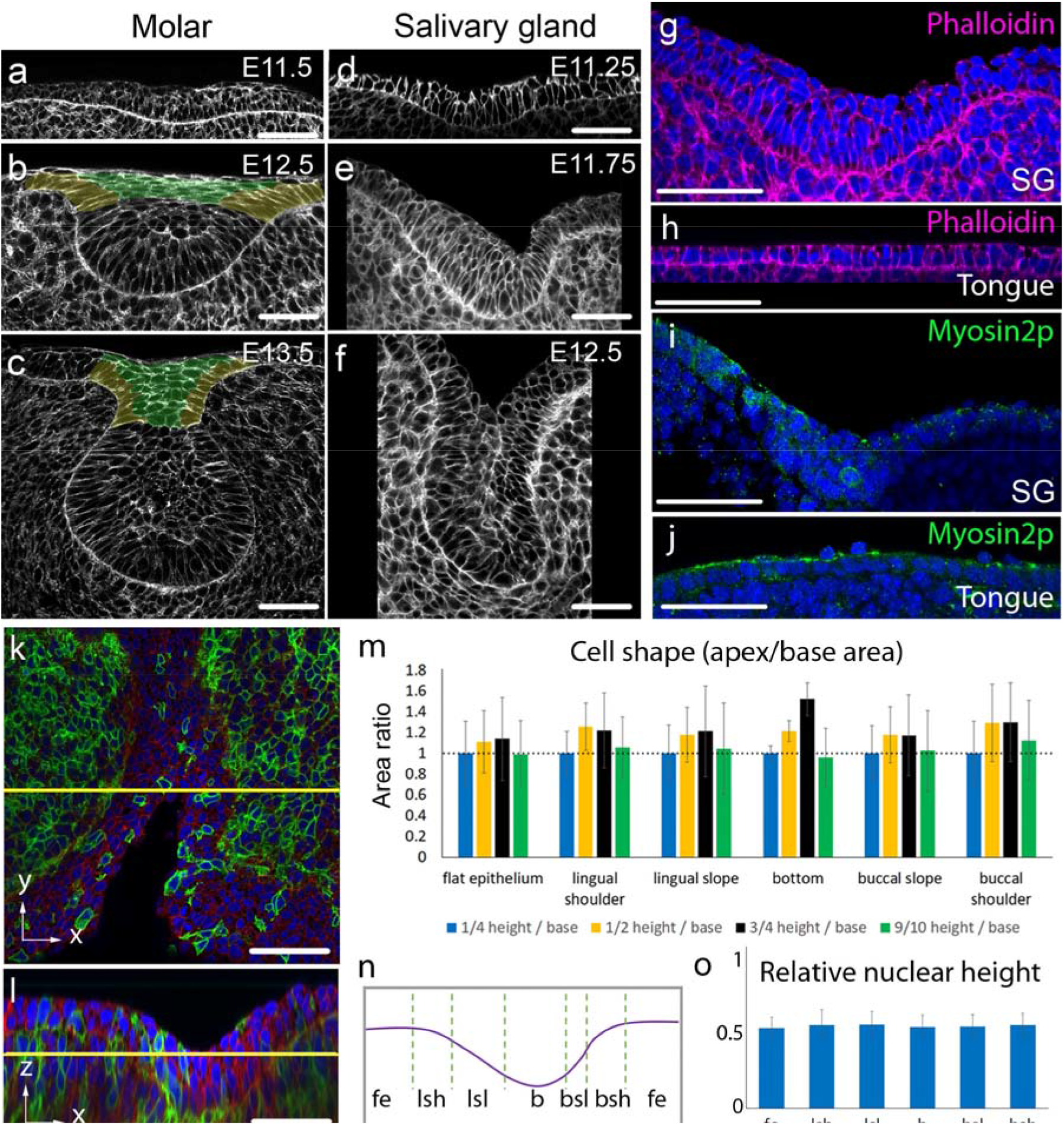
Invagination of early salivary placode is not driven by cell wedging. **a-f**: Phalloidin staining of indicated stages of molar tooth (a-c) and submandibular salivary gland (d-f). False-coloring indicates intercalating suprabasal cells (green) and shoulder/basal cells that transmit the contractile force to the basal lamina (yellow). **g-j:** Phalloidin (magenta), DAPI (blue) staining of the SG placode (g), tongue epithelium (h), and phosho-Thr18/Ser19 Myosin 2 immunofluorescence (green) of the SG (i), and tongue epithelium (j). **k-l:** Mosaically labelled mT/mG tissue stained by DAPI (blue) for cell shape and nuclear height analysis in SG. *En face* (k) and frontal (l) views. **m:** Ratios of planar cross-sectional areas of cells at indicated heights in different regions of the SG invagination revealing no wedging (3 litters, 11 placodes in total; p > 0.05 (one-way ANOVA)). **n:** Schematic showing regions across a salivary placode mediolaterally used in M and O. Fe: flat epithelium. Lsh: lingual shoulder region. Lsl: lingual slope. B: bottom. Bsl: buccal slope. Bsh: buccal shoulder. **o:** Nuclear position (height above lamina) as fraction of total cell height in indicated regions (3 litters, 4 placodes in total; p > 0.05 (one-way ANOVA)). Scale bars in all panels: 50 μm. Bar graphs are means +/− SDs.

Geometrically, the only way of packing a field of approximately cylindrical cells on an invaginated surface is by a staggered arrangement illustrated in Fig.2a. For this to be an active mechanism of invagination, the cells must undergo vertical shear, similar to the sliding between the steps of an escalator or sections of a “telescopic” arm. We therefore designated this cell rearrangement as “vertical telescoping”. In transverse sections of early SG we indeed observed numbers of vertically aligned cells arranged on the inclined, invaginating basal lamina (Fig.2c). To test whether vertical telescoping occurs throughout the invaginating SG in an unbiased way, we computationally segmented 3D confocal image stacks to determine the outlines of all the epithelial cells in 3D and fitted ellipsoids to every cell ^14^. Vertical telescoping definitively predicts that cells will lean or tilt outwards from the centre of the placode relative to the normal to the lamina (as shown schematically in Fig. 2a). We therefore mapped the angle of the ellipsoids’ principal axes to the plane of the immediately underlying basal lamina (Fig.2b). Measuring angles was preferred over attempting to measure vertical position differences due to the variability of cell heights. In early SG, we indeed observed a consistent outward lean between the cells and the lamina (Fig. 2e-g) as arrows pointing outwards radially from the placodal centre), unambiguously indicative of vertical telescoping across an entire placode. Judged by the length of the arrows, the extent of telescoping appeared correlated to the degree of invagination, i.e. the steeper the slope is, the more cells leaned outward from the placodal centre and quantification proved this to be the case (Extended Data Fig. 1, Table 1). Thus, vertical telescoping occurs throughout the SG.

**Fig. 2.**
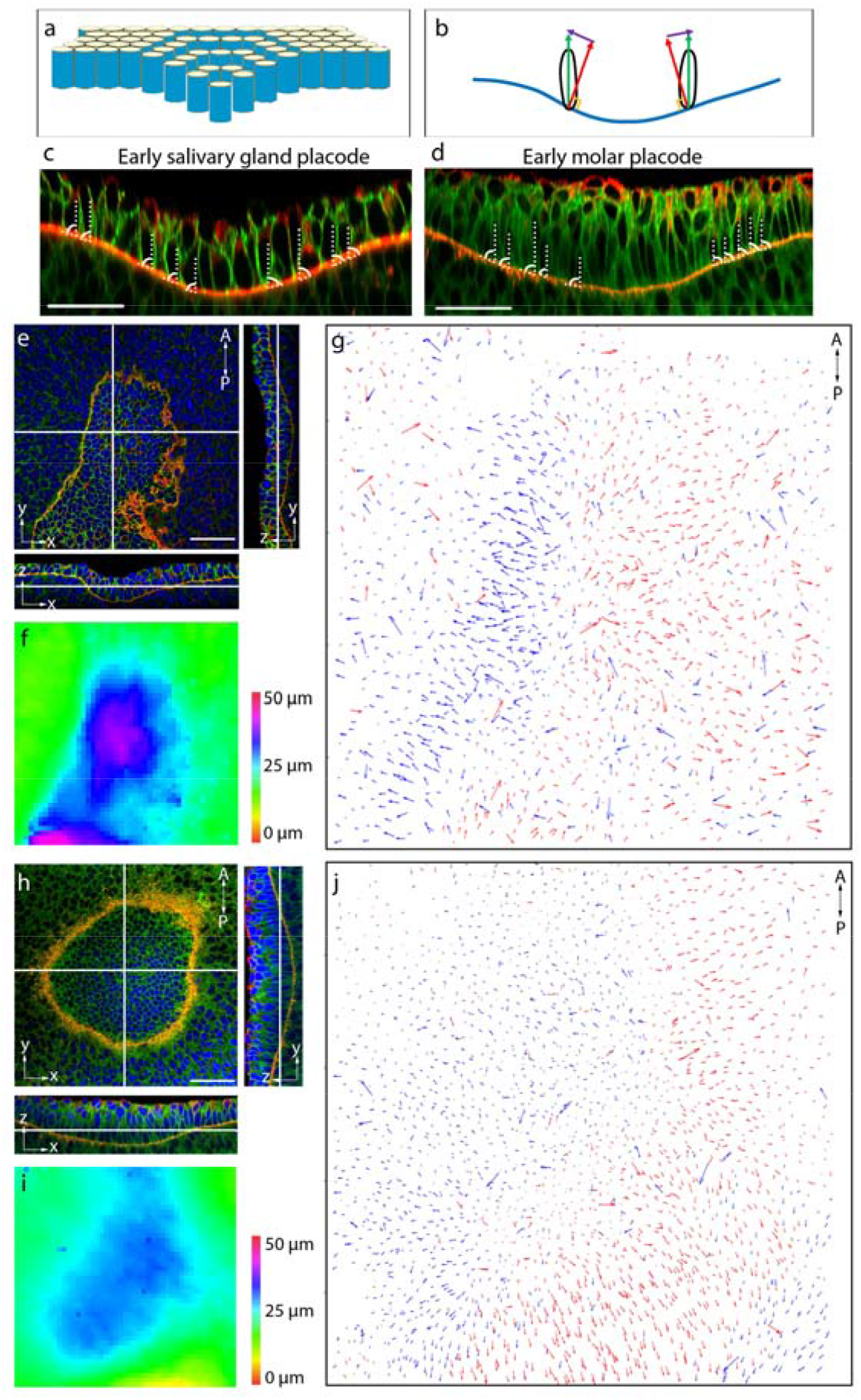
Vertical telescoping in early SG and molar invagination. **a**. Schematic showing vertical sliding between cells to generate invagination. **b**. Schematic illustrating the method of measuring and displaying cell tilt. Green vector: major axis of a basal cell. Red vector: normal vector of the local lamina. Purple vector: the displacement between the green and red vectors. Blue line: basal lamina of the placode. **c-d**. Immunofluorescence of transverse sections of SG and molar placode. Green: β-catenin. Red: Laminin-α1. Dashed lines: major axes of cells. Curved lines: the angle between the local lamina and the major axes of cells. **e** and **h**: Immunofluorescence of SG placode (**e**) and molar placode (**h**), top (xy plane) view and orthogonal (yz and zx plane) views. Green: β-catenin. Red: Laminin α1. Blue: DAPI. **f** and **i**: Heat map of the depth of the SG (**f**) and molar placode (**i**). **g** and **j**: Vector map of the displacement angle between cell and lamina vectors of the SG placode in **e** and the molar placode in **h**, indicating VT. Arrows are coloured by their direction to the left (blue) and right (red) to facilitate visualization. Note that due to the asymmetry of the SG placode, arrows on the left side (buccal side) of the placode are longer than those on the right (lingual), corresponding to the steepness of the slope. Maps are representative of 3 independent litters, for placode type and of 6 and 3 different placodes for SG and molars respectively. Scale bars: 50 μm

We then considered other ectodermal organs. Although different in later stages of development, the initial tooth primordium resembles early SG placode morphologically: both form an invaginated monolayer (before pseudostratification and then outright stratification by vertical cell divisions in the molar ^3, 15^). To test for vertical telescoping in the tooth, we examined molar primordia at their early initiation stages, when stratification has barely begun. Transverse sections revealed vertical cells on the inclined slope of the lamina (Fig. 2d). Mapping the cell-to-lamina angles in 3D, we saw that cells in the basal layer showed the tell-tale outward lean indicative of vertical telescoping, although with a central region having less-tilted cells, consistent with the flatter shape of the tooth invagination at its centre (Fig. 2h-j). Thus, vertical telescoping occurs in tooth primordia in the early stages of invagination before formation of a substantial contractile canopy. This shows a unity of morphogenetic mechanism between different placodal organs.

To understand the mechanism of vertical telescoping, we first determined whether it requires the underlying mesenchyme. If mesenchyme contributes to invagination, its removal should result in less invagination. Enzymatic removal of the mesenchyme resulted in a more, rather than less, invaginated placode (Extended Data Fig. 2). This result clearly shows that, rather than driving SG invagination, mesenchyme limits it at this stage and that there is most likely an epithelially autonomous mechanism for vertical telescoping.

If vertical telescoping is epithelially driven, then it implies active cell movement consisting of vertical cell migration of cells relative to their neighbours – in effect a “mesenchymoid” behaviour. By analogy with migration of cells on substrates, one might expect that the cells move with a leading-edge protrusion ^16^. For the vertical movement implicit in vertical telescoping, a leading edge could be either apical or basal (Fig. 3a, b), although some sort of novel snake-like undulating lateral movement is also theoretically possible (Fig. 3c) ^7^. Live imaging of mosaically labelled specimens revealed no apparent basal protrusions or lateral undulations, but did reveal rather highly conspicuous apical protrusions (Movie S1-S3). These protrusions were dynamic and somewhat less obvious, although still visible, in fixed material and in individual movie frames (Fig. 3d-e). Although some apical protrusions could be seen in inter-placodal flat epithelium (Extended Data Fig. 3a), those in the invaginating SG were much more abundant, much larger, and less static (Fig. 3f, Extended Data Fig. 3b, Movie S4 versus S1). Importantly, the protrusions in the SG cells were all directed towards its centre (i.e. centripetally) compared to the random orientation of non-placodal cell protrusions (Fig. 3g-j) (p < 0.001 inside placode and p > 0.05 outside placode, Rayleigh test). Thus, the protrusions are consistent with the model depicted in Fig. 3a: an apical centripetal leading edge somehow vertically raises a cell relative to its more central neighbour while depressing the latter to drive vertical telescoping.

**Fig. 3.**
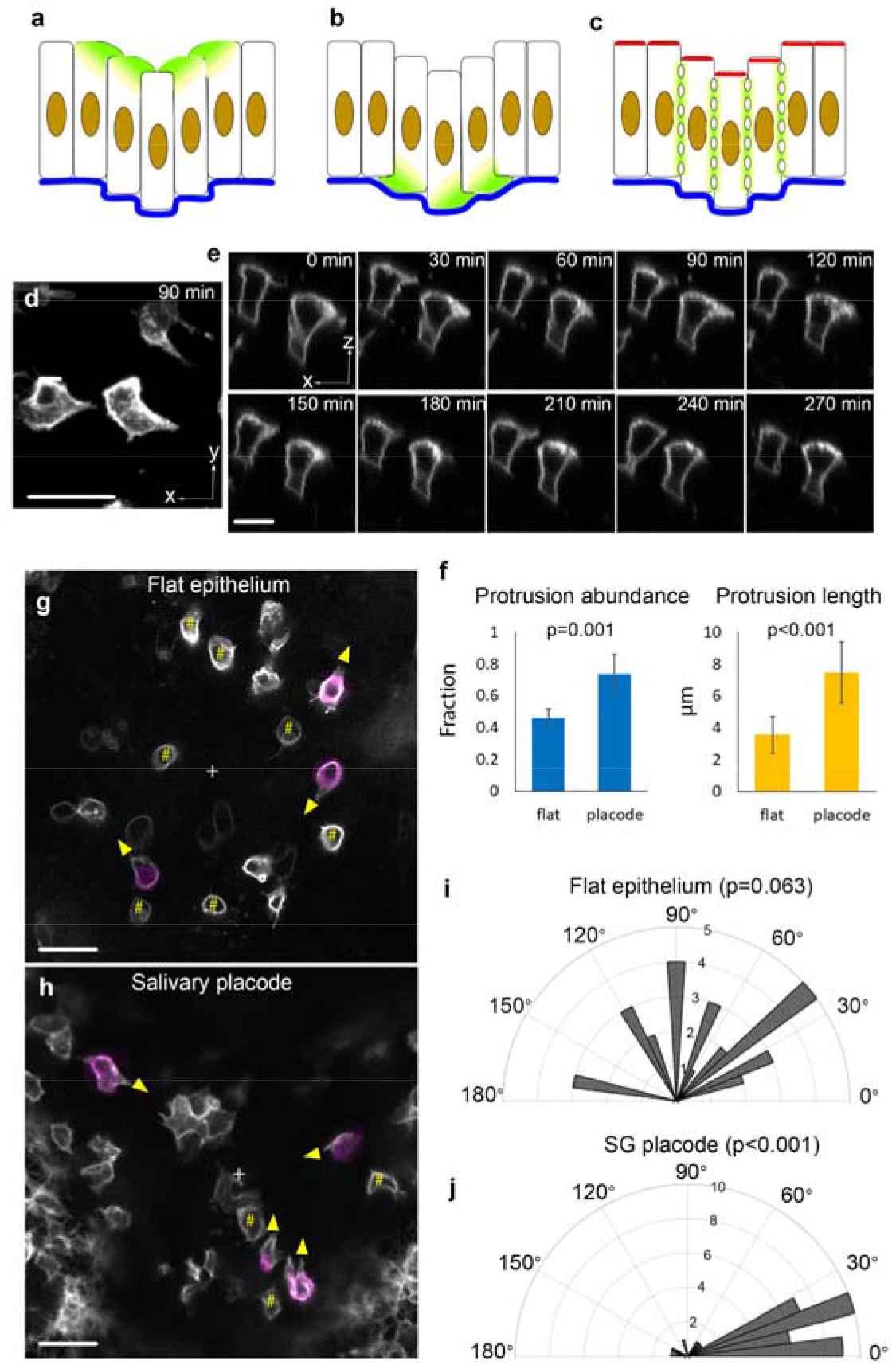
Centripetal apical protrusions are associated with VT. **a-c**: Schematics of possible cell behaviour in VT: apical protrusion (**a**), basal protrusion (**b**), and lateral undulation (**c**). **d**,**e**,**g**,**h**: Stills from movies of mosaically labelled epithelia. **d**: Top view of apical protrusions in SG placode from Movie S3 (placode centre is to the right). **e**: Side view of apical protrusions from Movie S2 (placode centre is to the right). Successive frames show protrusions are dynamic. **f**: Quantifications of the abundance and length of the protrusions. **g-h**: Top view of an area of flat epithelium (**g**) and SG placode (**h**) to show the orientation of apical protrusions (grayscale cell contours at the apical end extending beyond magenta-coloured cell contour at mid-height, yellow arrowheads show direction). Hash marks (#) indicate non-protrusive cells. “+” indicates centre of microscope field (**g**) or of placode (**h**). To avoid cell shape ambiguity, only isolatedly labelled basal cells were analysed. **i-j**: Quantification of orientation of protrusions (0° indicates protrusion towards field/placode centre. P values are calculated using the Rayleigh test for circular data (test for non-uniform disribution) and two-tailed unpaired t test for abundance and length data. Sample sizes: 3 independent litters, 6 SG placodes, 40 protrusive cells in total, 6 flat regions, 28 cells in total. Bar graphs are means +/− SDs. Scale bars: 20 μm.

To determine the function of the apical protrusions and signals that induce and orient them, we applied small molecule inhibitors to cultured SG placode explants. We titrated the concentrations of the actin polymerization inhibitor cytochalasin D and the Arp2/3 actin-branching inhibitor CK666 to the lowest concentrations that had a phenotypic effect. At these levels of cytochalasin D, the actin cytoskeleton is affected (Extended Data Fig. 4a) but cells remain remarkably intact (Extended Data Fig. 4c) and explants survive without shedding dead cells after 6 hours of incubation. CK666 has an even milder effect, with the actin distribution and cell shapes appearing broadly normal (Extended Data Fig. 4b,c). Both treatments caused a reduction in the number of apical protrusions (Fig. 4c) and inhibited invagination (Fig 4a,b). Given the selectivity of Arp2/3 for branched actin structures and their association with leading-edge cell protrusions, we conclude that the apical protrusions are likely to be important for the epithelial invagination.

**Fig. 4.**
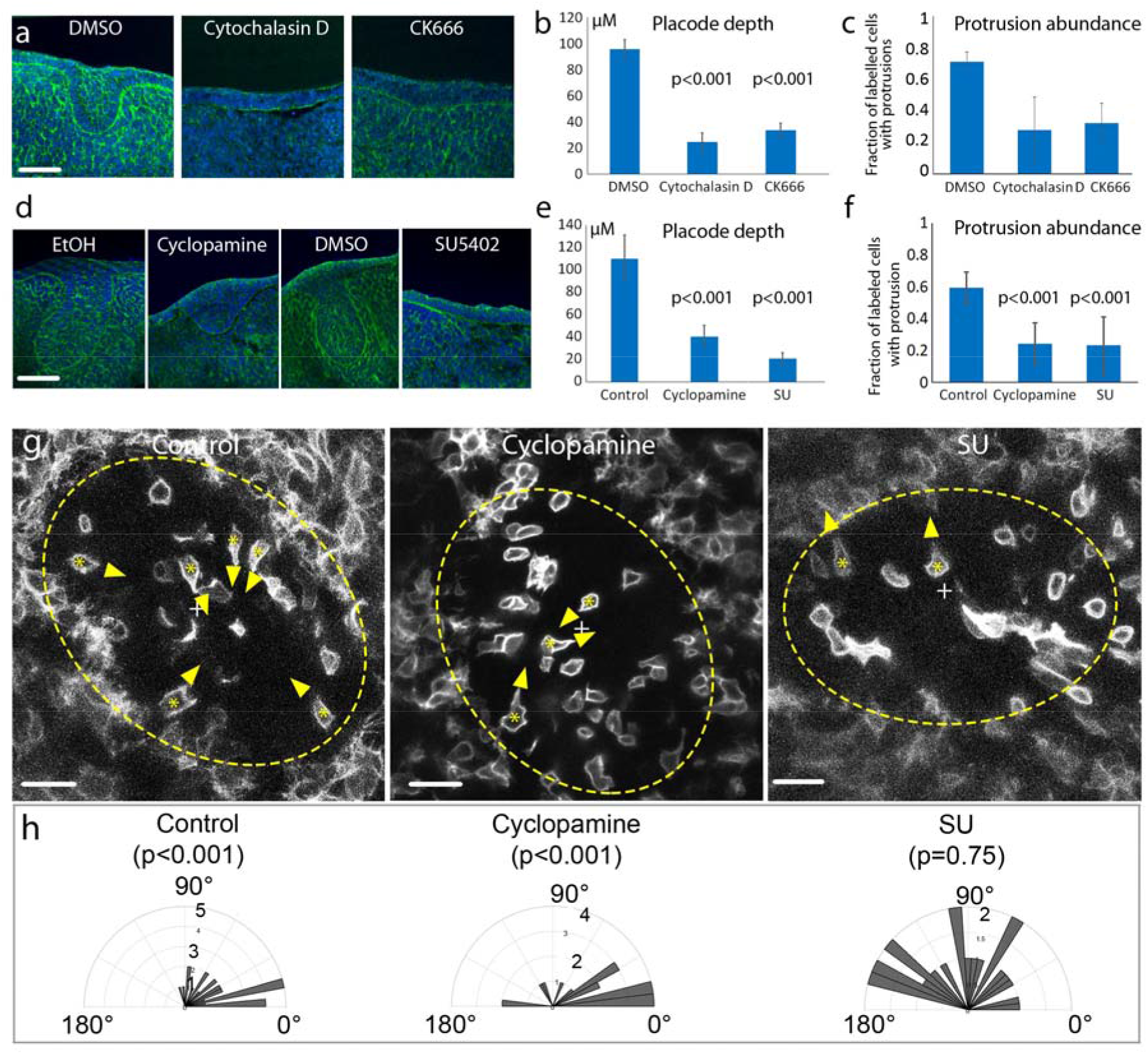
Protrusions and invagination are similarly sensitive to inhibitors. **a, d**: Transverse sections of SG placode explant culture with DMSO, Cytochalasin D, CK666, SU5402, ethanol, or cyclopamine as indicated (green: phalloidin; blue: DAPI) showing inhibition of invagination. Scale bars: 50 μm. **b, c**, **e, f**: Quantification of placode depth (b, e) and protrusion abundance (c, f) in inhibitor-treated placodes versus vehicle-treated controls (means +/− SDs from 3 independent litters for each treatment, 3 or 4 placodes per experiment per treatment). **c, f:** Quantification of by inhibitors compared to controls **g**. Top-view live images of SG placodes in mT/mG explants treated with vehicle, cyclopamine or SU5402. Dashed line circles: placode region. Asterisks inside cells: the cell bodies of protrusive cells. Arrowheads: direction of the protrusion. “+”: the centre of the placode. Note that protrusions are centripetal in control and cyclopamine treated tissues, but not in SU5402-treated tissue. Only isolatedly-labelled cells were analysed. Scale bars: 20 μm. **h** Orientation of protrusions in the three treatments (0° indicates centripetal) (data from 3 independent litters, 6 placodes in each condition). P values in **b, d**, and **f** are two tailed t-test, in **g** are Rayleigh test.

We previously showed ^15^ that both the Sonic Hedgehog (Shh) and Fibroblast Growth Factor (FGF) pathways drive early tooth morphogenesis. We therefore tested the effect of a Hedgehog signalling inhibitor (cyclopamine) and an inhibitor of FGF receptor (SU5402) in SG explants. At minimum concentrations known to inhibit expression of pathways targets (data not shown) cyclopamine reduced invagination and SU5402 essentially abolished it (Fig. 4d,e). Analysis of apical protrusions in mosaically labelled live tissue showed that both cyclopamine and SU reduced the number of protrusions (Fig. 4e-f). FGF pathway inhibition, unlike cyclopamine, also disorientated the protrusions from centripetal to random (Fig. 4g,h), potentially accounting for the complete loss of invagination and suggesting that FGF plays a polarizing or chemotactic role in organizing cell behaviour in vertical telescoping.

Together our findings reveal cell-on-cell migration as a previously unrecognized mechanism for epithelial monolayer invagination. All previously described and theoretical mechanisms going back at least to the 1930s and up to the present day, have involved cell wedging, while vertical telescoping does not. Thus vertical telescoping introduces a new principle of epithelial bending by cell rearrangement. It requires coordinated vertical migration behaviour of an ensemble of cells and could be considered a form of collective cell migration. Why has this mechanism not been described before? Possibly it does not occur in Drosophila, currently the premier system for epithelial morphogenesis. Possibly, it does not occur in early vertebrate development, although a re-examination of some instances of neural invagination may reveal it. We have shown vertical telescoping to occur in salivary gland and tooth, which are representative of the whole class of ectodermal organs. These are diverse, numerous, widespread and largely conserved across mammals and to some extent birds. Thus vertical telescoping is unlikely to be an obscure rarity.

The comparing the salivary gland with the tooth illustrates how vertical telescoping is related to canopy contraction ^3^. In both, cells migrate upwards and centripetally. In the tooth, but not the salivary gland, vertical cell division simply liberates apical (suprabasal) daughters, allowing them to migrate even more centripetally. They remain mechanically linked to their basal sisters, thus creating a contractile canopy coupled to the basal layer. The simple sum of the migratory and oriented cell division behaviours achieves the difference between these two types of ectodermal organ (Extended Data Fig. 5). The mechanics of vertical telescoping, as distinct from the kinematic that we describe, and the adhesive and cytoskeletal structures that are involved, remain to be established: mechanical modelling and experimental measurement of forces in this system is a current focus of further investigation.

Understanding mechanisms of physical morphogenesis is a necessary basis for improving the fidelity of organoids to their natural counterparts and for precision regeneration in vivo. Vertical telescoping is one such mechanism, a new type of fold in the art of epithelial origami, and a mechanistic module that can be combined with others to generate phenotypic diversity.

## Acknowledgments

We thank Eleni Panousopoulou and Ana Rolo for helpful discussions. The project was supported by BBSRC grant BB/P007325/1 to J.B.A.G.

## Author contributions

J.L performed experiments and data analysis; J.L and A.E developed computer pipelines and codes for image analysis; J.B.A.G. designed and supervised the project. All wrote the manuscript together.

**Materials and code requests & Correspondence** should be addressed to J.B.A.G.

## Materials and Methods

### Mouse strain and staging of embryos

All animal experiments were conducted under license from the United Kingdom Home Office and approval of the institutional Ethical Review Board. Live imaging of mosaically labelled cells, cell shape analyses in fixed materials, were performed using mT/mG mice [Gt(ROSA)26Sortm4(ACTB-tdTomato,-EGFP)Luo/J (Jackson Laboratories strain 007576)] crossed with a tamoxifen-inducible Cre male B6.Cg-Tg(CAG-cre/Esr1)5Amc/J (Jackson Laboratories strain 004682) ^3^. To induce mosaic expression of membrane GFP, 2 mg/animal Tamoxifen (Sigma) in corn oil was injected I.P. 24 hrs. prior to the experiment into pregnant mT/mG females, together with 1 mg progesterone (Sigma) to reduce fetal resorption. Automated cell orientation analysis, immunofluorescent staining, and inhibitor treatment were performed with CD-1 mice. Embryos were staged by nominal age (noon of the day of vaginal plug detection taken as E0.5).

### Whole-mount immunostaining and imaging

Specimens were fixed in 4% paraformaldehyde for 3 hrs. at room temperature (RT). Fixed whole-mount tissues were washed 3 times with PBS-Triton (0.1%) and blocked with 20% goat serum (Sigma G6767) for 1 hr. at RT. Antibody and chemical staining were performed in blocking solution at the following dilutions: anti-β-catenin 1:500 (Sigma C2206), anti-laminin 1:500 (Sigma L9393), anti-phosphorylated myosin 2 light chain (Cell Signaling 3674), Alexa 488/568 anti-mouse/rabbit/rat 1:500 (Life Technologies), fluorescent phalloidin (Life Technologies) 1:200, DAPI (Life Technologies) 1:10000. Validations of antibodies can be found on manufacturers’ websites. Incubations for both primary and secondary antibodies were at 4°C overnight. For specimens >80 μm thick, post-fixation (4% PFA, RT, 1 hr.) was performed after immunostaining, and stained tissues were cleared using Scale A2 and Scale B4 solutions ^17^ for confocal imaging.

3D image stacks were captured on a Leica SP5 confocal microscope with an HCX PL APO CS 40X oil (N.A. 1.25) or an HCX PL APO 63x / 1.3 GLYC CORR CS (21° C) objective. For 3D cell shape and orientation analyses, placodes were imaged *en face* at optimal z resolution.

### Cell shape and nuclear height analyses

For cell shape analysis, GFP positive embryos were dissected and submandibular glands were imaged *en face* at optimal z resolution under the 63X objective. Isolated GFP-positive cells contacting the basal lamina were identified by digitally reslicing the 3D stack frontally and viewing in frontal section. To measure cell shapes, along the apicobasal axis, the apical and basal limits of a cell were decided by scrolling up and down the original image stack to find where in z the GFP cell boundary started and ended. These two positions were marked and slice numbers of a quarter, half, three quarters and 9/10 cell height from the base were calculated. Areas of the cell inside the boundary at each of these heights were drawn manually using the freehand tool in Fiji to return the area enclosed. “Base” area for Fig. 1Q was taken as that of the quarter-height slice (necessary because the actual base is often oblique) and the “apical” area was that of the 9/10-height slice (for similar reasons). For nuclear height analysis, whole mount mandibles were stained with DAPI to show the nucleus. Submandibular glands were imaged *en face* as 3D stacks at optimal z resolution under a 63X confocal objective. The centre of the nucleus and the height of each cell were decided by viewing the cell on a transverse digital section that shows the largest section area of the cell. The line tool in Fiji was used to manually measure the height of the nucleus and the cell. ANOVA test was performed to decide whether there is any significant difference between groups.

### Automated quantification of cell orientation

Whole-mount mandibles were stained for β-catenin (for cell boundary), α-laminin (for basal lamina), and DAPI (for nucleus), and a 3D stack of the placode was imaged *en face* at optimal z resolution on a Leica SP5 40x or 63x objective. Fiji macro and MATLAB were used to segment epithelial cells in 3D and compute cell orientations, respectively (scripts available upon request). Essential steps in the pipeline are as follows: an entire image was divided in x-y as a grid (each square 20 × 20 pixels), keeping the z depth. The 3D coordinates of the local lamina were registered as the peak of the laminin staining intensity in z and the x-y coordinates of the small square. The mesenchyme was blacked out from the image below the lamina. Cells were segmented in 3D using our self-developed iterative thresholding Fiji macro and the 3D suite Fiji plugin ^14^ based on the cell boundary staining, and the segmented objects were filtered for DAPI staining to confirm them as real cells. Ellipsoids were fitted to each segmented cell volume using the Fiji ellipsoid-fitting macro ^14^ and information of cell centroids and axes of the ellipsoids were extracted. These cell data together with the lamina grid positions were imported into MATLAB. The plane of the local lamina of a cell was determined as that defined by the xyz coordinates of the lamina at the centre points of the three 20×20 pixel regions closest to the centroid of the cell. Orientation of a cell was computed in MATLAB as the angle between the major axis (as a vector) of the cell and the normal vector to the local lamina plane. Displacements of these two vectors were plotted as a vector map to show the “leaning” of columnar cells on a bending lamina. A map of the depth of the basal lamina was plotted as a heat map in R ^18^. For scatter plots in Extended data Fig. 2, cells were defined as basal if their basal-most points were ≤ 3 μm above the basal lamina plane. Code for the above procedures is available on request from the authors.

### Explant Culture and treatments

Explant culture of salivary placodes was performed as described in ^15^. Concentrations of drugs used in culture were: 20 μM cyclopamine (Sigma, diluted from 10 mM stock in ethanol), 5 μg/ml SU5402 (Sigma, diluted from 10 mg/ml stock in DMSO) 0.05 μg/ml cytochalasin D (Sigma, diluted from 10 mM stock in DMSO), and 250 μM CK666 (Sigma, diluted from 80 mg/ml stock in DMSO). Ethanol or DMSO were used as vehicle, respectively. Culture medium is phenol red-free Dulbecco’s Modified Eagle Medium: Nutrient Mixture F-12 (DMEM/F12) + 15% FCS + 0.1g/L Ascorbic Acid + Pen/Strep.

Sample sizes of each experiment are detailed in each figure legend.

### Live imaging

GFP-positive mT/mG embryos were selected for live imaging. Submandibular gland explants were dissected at E11.25 in DMEM/F12, immobilized apical-side-down in a glass bottom dish by overlaying with a piece of transparent membrane from a porous cell culture insert (Corning 353090, 0.4μm) anchored to the glass at its edges with small blobs of Vaseline, and imaged on an inverted Leica SP5 with a 63X objective with optimal z resolution. The culture was supplied with humidified 5% CO_2_ in air in a 37°C incubation chamber throughout the imaging.

To assess the effect of drugs on cell protrusions in live tissue, mandibular explants were dissected and cultured with drugs in concentrations listed above, in a glass bottom dish and shallow medium so that the surface of the tissue was in contact with air (for tissue health). Explants were cultured at least 3 hrs. before setting up for imaging.

