## Supplemental Information for "Epithelial invagination by vertical telescoping"

Extended Data

**This file includes:**

Extended data Figs.1 to 5

Table S1

Captions for Movies S1 to S5

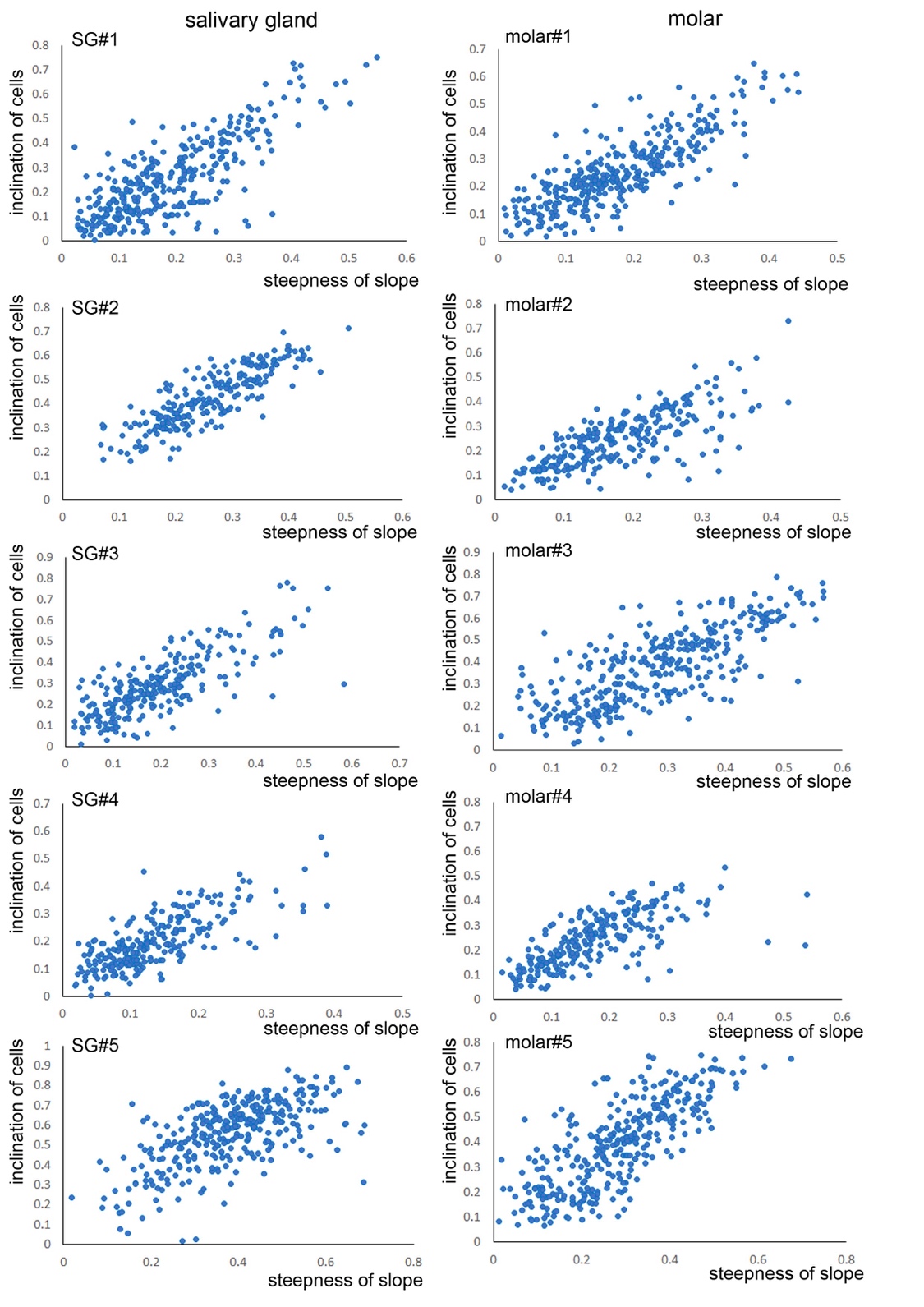

Extended Data Fig. 1. The inclination of epithelial cells is correlated with the steepness of the placodal slope. Amount of deviation of the cell axis from normal (perpendicular) to the basal lamina is plotted against the lamina angle-to-the-horizontal. Both are expressed as the cosine of the angle.

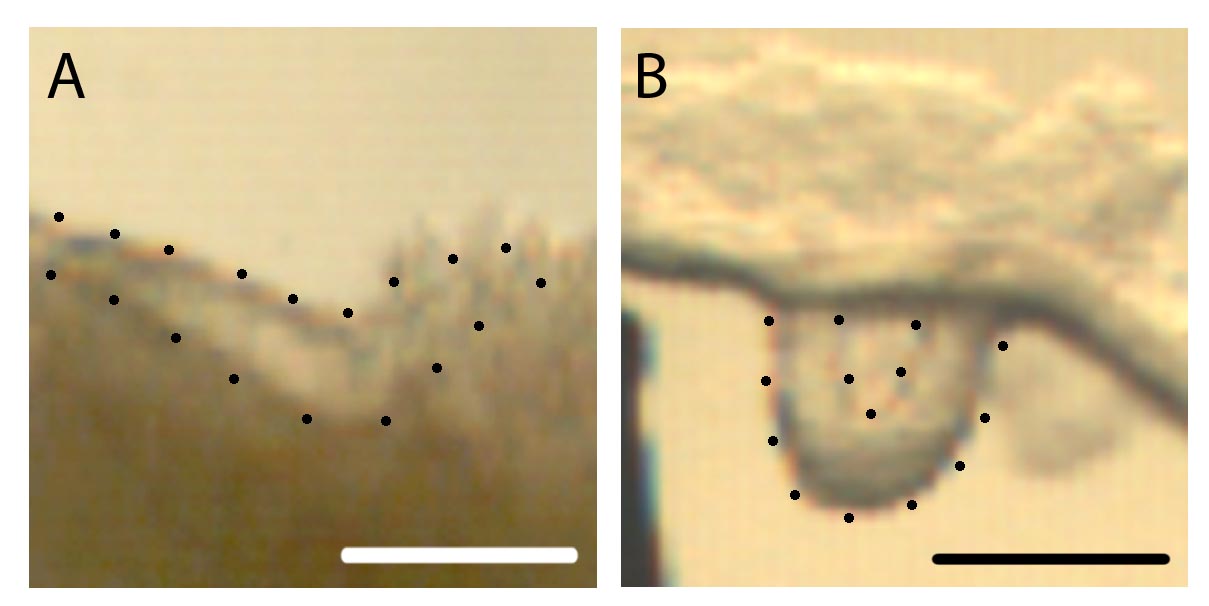

**Extended Data Fig. 2. SG epithelium invagination is deeper upon enzymatic removal of underlying mesenchyme: a**. Bright field image of an intact E11.5 salivary gland placode explant (frontal slice) with its underlying mesenchyme showing modest v-shaped invagination of the epithelium (apical and basal surfaces indicated by dotted lines) consistent with this stage.

**b**. Bright field image of an identically-staged littermate salivary gland placode enzymatically dissociated from the mesenchyme showing much deeper invagination.

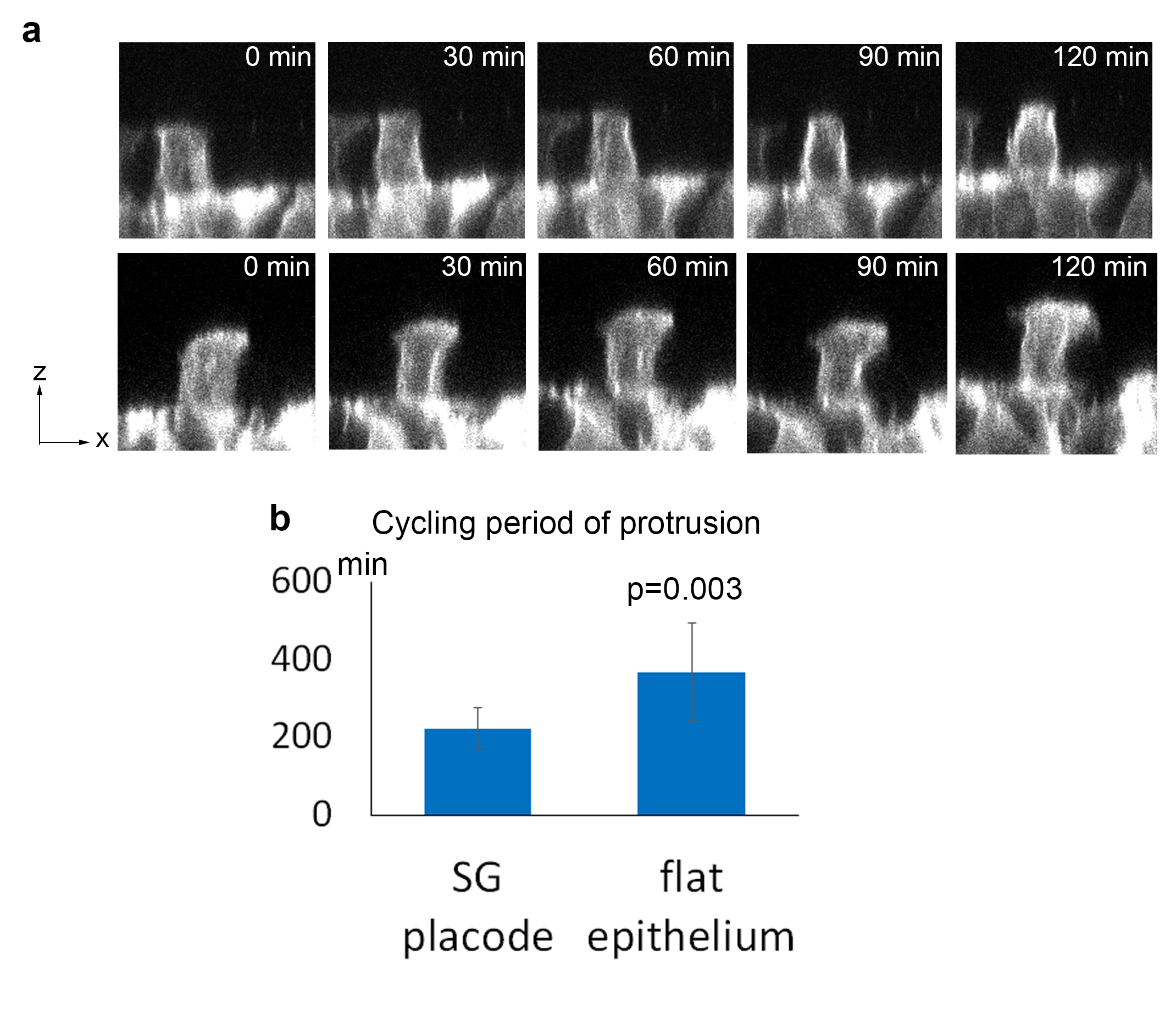

Extended Data Fig. 4. Apical planar protrusions are active in placode epithelium

**a**. Example stills from live imaging of a flat epithelial region. Top panels: a non-protrusive cell. Bottom panels: a protrusive cell. **b**. Measurement of the persistence period of individual protrusions in cells in flat or invaginating regions. Graph shows means and SDs. P value: t-test. Related to Movie S4.

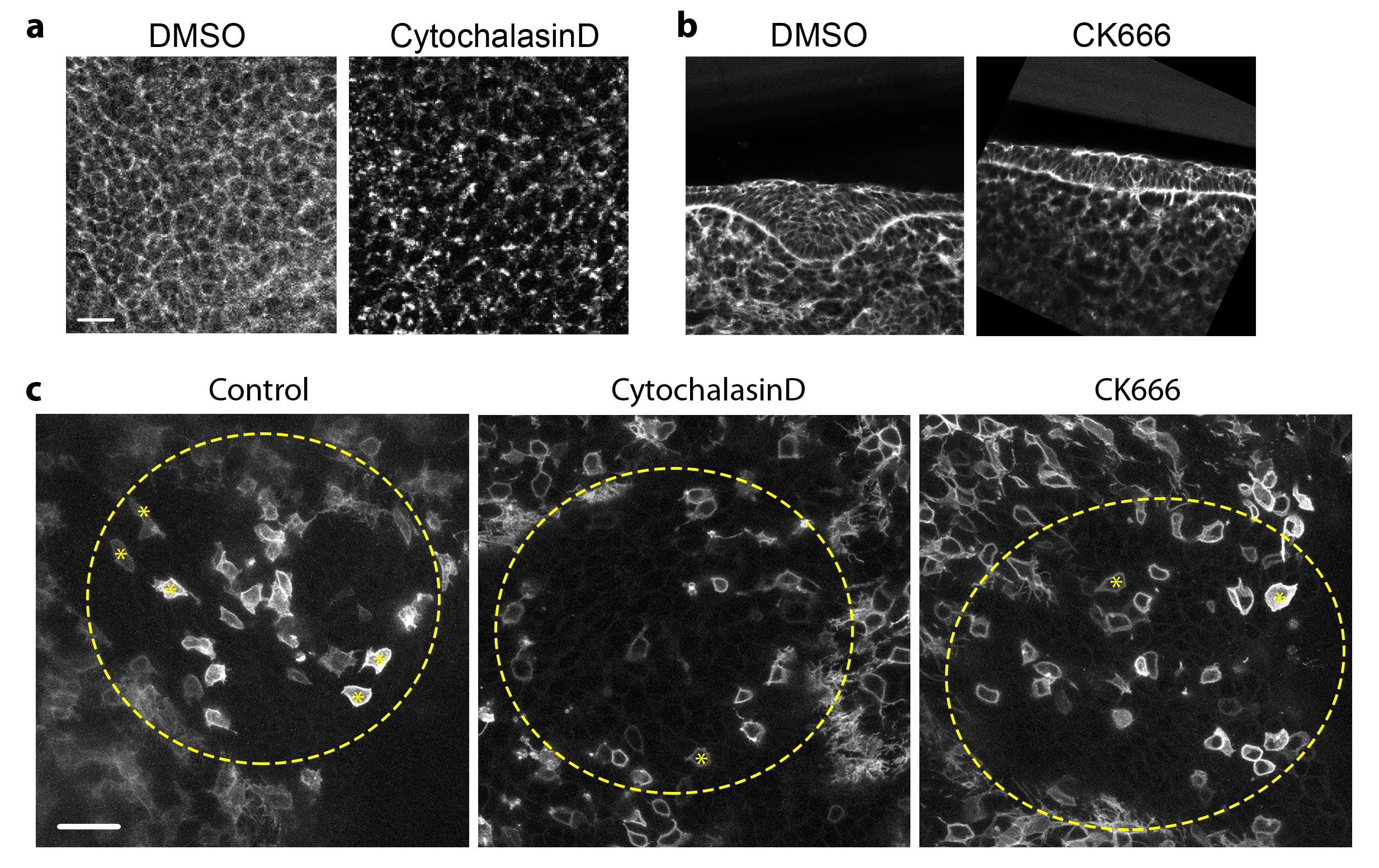

**Extended Data Fig. 5. Effect of cytochalasin D and Arp2/3 inhibitor CK666 on F actin and protrusions.** **a**. Phalloidin staining of the epithelium cultured with vehicle (DMSO), cytochalasin D (top views). **b**. Phalloidin staining of mandible slice cultured with vehicle (DMSO) and CK666. **c**. *En face* live images of SG placode of an mT/mG tissue treated with control vehicle, cytochalasin D or CK666. Dashed line circles: placode region. Asterisks inside cells: the cell bodies of protrusive cells. Cross: the centre of the placode. **d**. Quantification of the abundance of protrusions in the three conditions. Statistics are based on 3 independent experiments, at least 6 placodes in total in each condition. Bar graphs are means and SDs. P values in **d** are two-tailed t test. Scale bars: 20 μm.

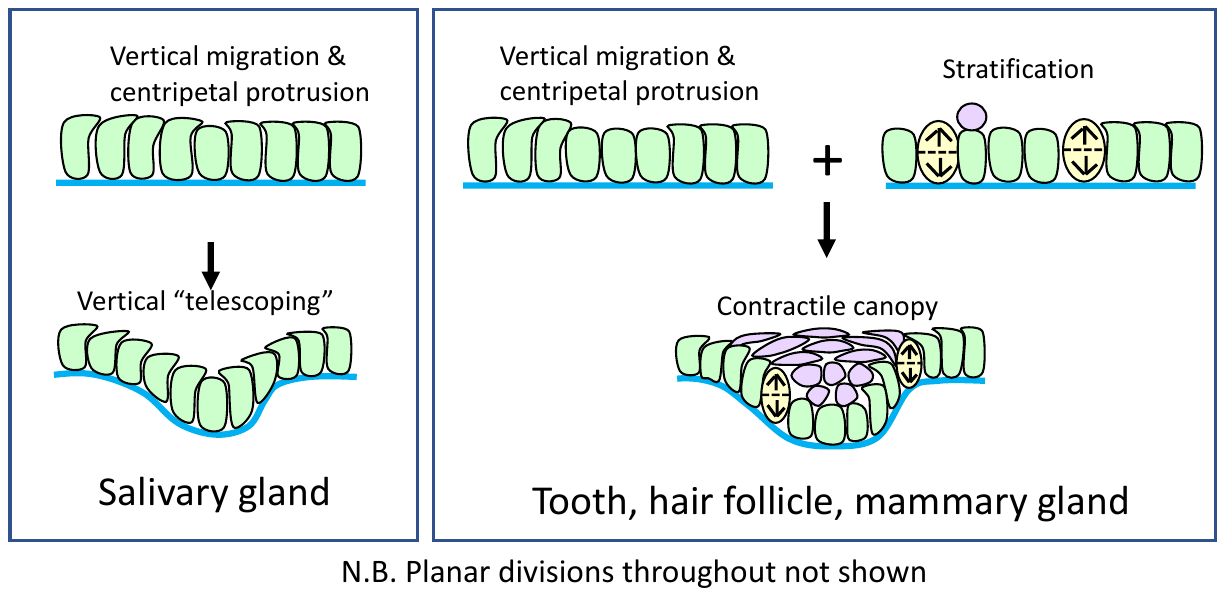

**Extended Data Fig. 6.** **Tooth morphogenesis is a composite of vertical telescoping plus canopy contraction.** Schematic of epithelial invagination in the SG and molar, respectively by VT alone or a simple combination of VT, vertical cell division, and canopy contraction by suprabasal cell intercalation. Blue line: basal lamina. Green: basal cells. Yellow: dividing cells. Lilac: suprabasal cells.

|  | Cell number | R-squared value |
| --- | --- | --- |
| SG#1 | 236 | 0.7134 |
| SG#2 | 346 | 0.6197 |
| SG#3 | 255 | 0.59 |
| SG#4 | 270 | 0.5298 |
| SG#5 | 334 | 0.4017 |
| Molar#1 | 357 | 0.6635 |
| Molar#2 | 262 | 0.5672 |
| Molar#3 | 333 | 0.579 |
| Molar#4 | 246 | 0.5052 |
| Molar#5 | 340 | 0.5964 |

Table S1. Number of cells and Correlation coefficients for cell inclination to lamina versus lamina slope.

**Movie S1. SG epithelial cells extend centripetal apical protrusions – side view**

Transverse view of mosaically GFP-positive cells in a SG placode, showing centripetally directed apical protrusions (arrowed). E11.25 mT/mG embryos were injected with Tamoxifen to generate mosaic labelling. Mandible explants were dissected from GFP positive embryos and mounted in DMEM/F12 medium to be imaged live *en face* as described in Materials and Methods. Frame interval: 30 min. 10 frames in total.

**Movie S2. SG epithelial cells extend centripetal apical protrusions – top view**

*En face* view of video stack in Movie S2 showing centripetal apical protrusions. Frame interval: 30 min. 10 frames in total.

**Movie S3. Apically protruding cell shape**

3D rendering of GFP-positive basal epithelial cells on a SG slope showing characteristic oblique base and large apical protrusion.

**Movie S4. A few non-placodal cells have small quiescent protrusions**

Side view (XZ digital slice) of a mosaically labeled mandible explant in a flat region of the epithelium showing two non-placodal cells (at top) with protrusion that are essentially non-motile compared to those in the control. Frame interval: 30 min. 12 frames in total.
